# Annotating RNA motifs in sequences and alignments

**DOI:** 10.1101/011197

**Authors:** Paul P. Gardner, Hisham Eldai

## Abstract

RNA performs a diverse array of important functions across all cellular life. These functions include important roles in translation, building translational machinery and maturing messenger RNA. More recent discoveries include the miRNAs and bacterial sRNAs that regulate gene expression, the thermosensors, riboswitches and other cis-regulatory elements that help prokaryotes sense their environment and eukaryotic piRNAs that suppress transposition. However, there can be a long period between the initial discovery of a RNA and determining its function. We present a bioinformatic approach to characterise RNA motifs, which are the central building blocks of RNA structure. These motifs can, in some instances, provide researchers with functional hypotheses for uncharacterised RNAs. Moreover, we introduce a new profile-based database of RNA motifs - RMfam - and illustrate its application for investigating the evolution and functional characterisation of RNA.

All the data and scripts associated with this work is available from: https://github.com/ppgardne/RMfam

## 1. Introduction

Characterising functional RNAs is an extraordinarily difficult task. Even highly transcribed RNAs from model organisms have remained uncharacterised for decades after their discovery. A specific example is the 6S sRNA, which was discovered in 1971. The 6S sRNA is conserved across Bacteria and is highly expressed in stationary-phase cells [1, 2]. But the role of 6S as a regulator of RNA polymerase remained an enigma for almost three decades [3]. Likewise, Y RNA, which was discovered in 1981, is broadly conserved across metazoans and is highly expressed [4]. It took two and a half decades before Y RNAs were shown to be essential for the initiation of DNA replication [5]. However, the mechanism for Y RNA function still remains unclear. These and similar examples show that it is remarkably difficult to functionally characterise RNAs, even after decades of work.

A new generation of tools for RNA discovery is now available thanks to powerful new sequencing technologies. Entire transcriptomes from species at different life stages, tissue types and conditions can be studied with RNA-seq [6, 7, 8]. The total complement of RNA structures encoded in transcriptomes is also accessible with SHAPE-seq [9] and functional regions of entire genomes of bacteria can be probed with techniques like TraDIS and Tn-seq [10, 11]. The data obtained by these tools are unearthing novel RNAs at an unprecedented rate, many of which are evolutionarily conserved, highly expressed, activated under specific conditions, essential and fold into conserved secondary structures. Annotation efforts such as those by the Rfam consortium [12, 13] are useful. However, many RNAs are not found in this database and many that have been curated remain uncharacterised [8]. To make sense of the volumes of transcriptome data that is now being generated, annotating this data and functionally characterising the cohort of RNAs of Unknown Function (RUFs) is critical. A complication for such work is that evolutionary turnover, as well as sequence variation can be high for ncRNAs [14, 15]. Consequently homology searches and other sequence-alignment based analyses can be very challenging.

Many RNAs contain functional structures that recur both within and across different RNA families. These motifs provide signatures that can identify functional components of RNA sequences. The motifs that have been characterised to date are involved in a diverse number of functions. These include imparting structural stability, facilitating interactions with other biomolecules, specifying cellular localisation and coordinating gene regulatory signals [16, 17, 18, 19]

A number of publications detail bioinformatic methods for the *de novo* discovery of RNA motifs from RNA primary sequences [20, 21]. There are also tools that can screen RNA secondary structures [22] and RNA tertiary structures [23]. The *de scito* (knowledge-based) approaches for the annotation of RNA motifs include sequence and structure descriptors [24, 25], primary and secondary structure-based profile methods for specific motifs [26, 27] and even methods that combine primary, secondary and tertiary data [19]. We complement these approaches by introducing a resource that identifies a range of previously characterised RNA motifs in RNA sequences and alignments using covariance models (CMs) [28, 29, 30, 31, 32].

We present 34 alignments, consensus structures and corresponding probabilistic models of published RNA motifs. We call this resource RMfam, or RNA Motif Families (all associated data and computer code is freely available from our repository hosted on GitHub: http://github.com/ppgardne/RMfam). These have been used to predict approximately 1, 900 conserved motifs in the Rfam (v11.0) alignments of RNA families; many of which are confirmed in the published literature. Finally, we show examples of the applicability of our approach for studying RNA function, evolution and alignment curation.

## 2. MATERIALS & METHODS

### 2.1 Distinction between Rfam and RMfam

The Rfam database collects and curates “seed alignments” of RNA families. These are non-coding RNAs, cis-regulatory elements and self-splicing introns. The alignments are manually constructed and annotated with consensus secondary structures, and used to seed probabilities for covariance models (CMs) for each family. The Rfam CMs are widely used for genome annotation projects to identify RNA loci (e.g. [33]). A requirement before each family can pass Rfam quality-control is that it is specific. In other words, there exists a bit score threshold for each CM that distinguishes between sequence matches that are related to the family and obvious false-positive matches. Consequently, many RNA motifs are not included in Rfam as they lack the required specificity [34, 35, 36, 12, 13].

### 2.2 What is an RNA motif?

For the purposes of this work an RNA motif is a non-trivial, recurring RNA sequence and/or secondary structure that can be predominantly described by local sequence and secondary structure elements. The motif is generally not restricted to a particular family or taxonomic group. Note that in other contexts, such as structural biology, a more general definition of motif is frequently used, e.g. [37].

Accurate probabilistic methods for annotating structured RNAs on DNA sequences called hidden Markov models (HMMs) and covariance models (CMs) are now available [28, 29, 30, 31, 32, 38]. From a given alignment, probabilistic models of conserved sequence (HMMs) and conserved sequence plus secondary-structure (CMs) can be built and used to filter large numbers of sequences for candidate homologous and/or analogous regions [39]. CMs cater to the characteristics of RNA sequence evolution that are imposed by basepairing (i.e. variation tends to preserve basepairing), the result is that the accuracy of CMs is greater than alternative approaches [40]. The computational speed of CMs has tended to be poor, however a lot of effort has been expended on improving the speed of the approach while maintaining the accuracy. Improvements include using HMMs as pre-filters to accelerate CMs, query-dependent banding and Dirichlet mixture priors [41, 39, 42, 38, 43].

**Figure 1.**
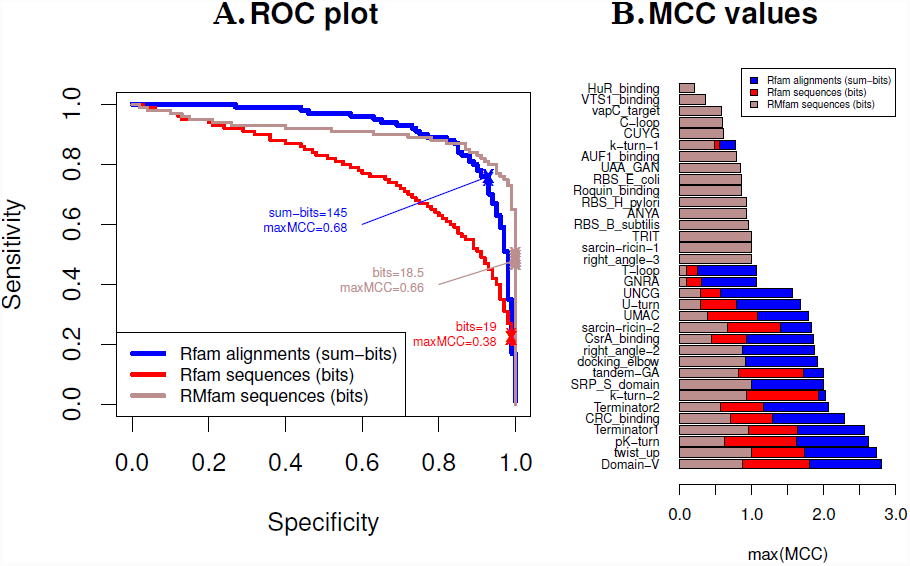
In the above plots we assess the accuracy of motif annotation and test whether annotating alignments instead of sequences improves the prediction accuracy. We have applied three different benchmarks (described in the text). In sub-figure **A** we show a ROC plot for pooled RMfam annotations. This plots the sensitivity versus specificity of all the motif annotations on sequences or alignments at different score thresholds. The ’x’s illustrate where on the curve the maximum Matthew’s Correlation Coefficient is located, and the corresponding bit scores are indicated. In sub-figure **B** we illustrate the maximum MCC of CM annotation for each motif from the 3 different benchmarks.

RMfam sequences, structures and alignments were collated from a variety of heterogeneous and sometimes overlapping data repositories [12, 23, 44, 27, 45, 46, 47, 48, 37, 49, 50, 51]. Where possible we sourced data from publicly accessible RNA motif resources, these included the FR3D MotifLibrary [37], the models supplied with RMDetect [19], the comparative RNA website [47] and SCOR [46]. We also used information from specialised resources such as the k-turn structural database [44] and SRPDB [52], as well as generating our own alignments for motifs such as the Shine-Dalgarno and Rho-independent terminators based upon the context of genome annotations (e.g. [27]). RNAFrabase was frequently used as a source of RNA secondary structure annotations derived from PDB structures [53, 54]. Finally, where necessary, we extracted sequences from publications. This was often a manual effort, involving manually transcribing sequences and structures from figures in published manuscripts. Where possible, these were mapped to PDB (downloaded June 2014) nucleotide sequences [55, 56, 57], the EMBL nucleotide archive [58] and Rfam (v11.0) [12, 13]. The provenance of each dataset is stored in the corresponding Stockholm alignment. Each of these motifs were then passed through quality control steps, where the sensitivity and specificity of the resulting motif is assessed (See Figures 1 and S10-S43). If these failed (e.g. the CM cannot identify member sequences or the false-positive rate is extremely high), then the motif was not included in the database. Each motif is also assigned a curated score-threshold. This threshold (in bits) provides a reasonable distinction between true and false matches.

### 2.3 Benchmarking motif annotations

In the following we briefly describe the benchmarks we have used to evaluate our motif annotations. These are described in further detail and with more elaborate results in the Supplementary Materials.

In order to determine the accuracy of our approach we ran a series of three benchmarks. These were evaluated on individual motifs (see Figures 1B and S10-S43), as well as on the collective RMfam results (see Figures 1A and S9). The first uses “RMfam sequences” which are taken from the seed alignments. Ten shuffled sequences, with identical dinucleotide distributions, were generated for each RMfam seed sequence [59]. Together these serve as positive and negative controls for our test.

We constructed two further tests based upon Rfam (v11.0) families. We identified Rfam families where there exists good evidence (primarily based upon literature) that a motif is conserved in the family of related sequences (Supplementary Table 1). These serve as positive controls for two further tests. For the “Rfam sequences” benchmark we randomly selected at least five sequences from each Rfam seed alignment (if fewer than five sequences were available, then all were included). We generated ten shuffled versions of each sequence; all had an identical di-nucleotide distribution to the native sequence. These sequences were all annotated with RMfam motifs, their CM scores were recorded and used to evaluate the accuracy of the annotations. Finally, for a “Rfam alignments” benchmark, we evaluated the accuracy of RMfam annotations in an alignment context. Each Rfam alignment was filtered, removing sequences more than 90% identical. The remaining sequences were annotated with RMfam CMs, retaining only those that cover more than 10% of the seed sequences and more than two Rfam seed sequences. The summary statistic we use for this final benchmark is a “sum-bits” score, this is the sum of the bit scores for each match in all the sequences in a seed.

The accuracy metrics that we report here are the Matthew’s correlation coefficient (MCC) [60], sensitivity and specificity.

The CMs built from RNA motifs tend to be short and contain little sequence information. In RMfam the mean sequence length is just 34.3 nucleotides and the mean number of basepairs is 10.9. Therefore scans of large sequence databases with these models result in a number of false-positives. We propose that annotating sequence alignments of ncRNAs have the potential to improve the specificity of our predictions. This assumes that evolutionarily conserved motifs are more likely to be correct. In theory this approach could be extended to genome alignments of e.g. transcribed regions.

## 3. RESULTS

In this study we present 34 RMfam alignments and probabilistic models of published RNA motifs (all freely available from our repository hosted on GitHub: http://github.com/ppgardne/RMfam). These have been used to predict approximately 2,500 conserved motifs in the Rfam (v11.0) seed alignments; many of which are confirmed in the published literature. Furthermore, our permutation tests have shown that both the sensitivity and specificity of this approach is remarkably high given the short motifs we use (See Figures 1 and S9-S44).

### 3.1 Function

One of the most labour intensive stages of RNA research is identifying the function of newly discovered RNAs. In order to illustrate the utility of RMfam for this task we show the matches between a model of the CsrA-binding site and two RNA families of unknown function, TwoAYGGAY and Bacillaceae-1 (Rfam IDs RF01731 and RF01690, see Figure 2). CsrA is a bacterial RNA binding protein that regulates the translation and stability of mRNAs [62]. It binds mRNAs carrying CsrA binding motifs, physically occluding ribosome-binding sites. This binding can itself be regulated by competition between the mRNAs and highly expressed sRNAs that host numerous CsrA binding sites. However, this class of sRNA (CsrB, CsrC, RsmX, RsmY and RsmZ) has only been identified in Gammaproteobacteria [63, 64]. The ncRNAs, TwoAYGGAY and Bacillaceae-1, were initially discovered in a large-scale bioinformatic screen [65]. Some further analysis identified two tandem-GAs in one of the stems that characterise the structure of TwoAYGGAY [19]. The matches between these families and the CsrA binding motif were discovered in this work and provide a testable hypothesis for further validation that there are also CsrA binding sRNAs in Clostridia (TwoAYGGAY), and Bacillales and Lactobacillales (Bacillaceae-1). The validation of these predictions is a work in progress with our collaborators.

**Figure 2.**
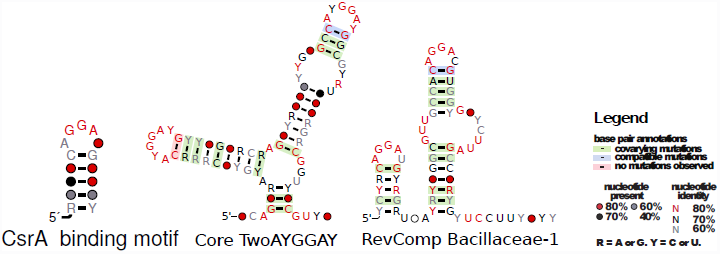
The secondary structures and sequence conservation of CsrA binding motif and two new candidate CsrA binding sRNAs, TwoAYGGAY and Bacillaceae-1 illustrated with R2R [61]. These families each have two strong matches to the CsrA-binding motif, this new evidence provides a strong case that these RNAs regulate the activity of the regulatory protein, CsrA, by sequestering this nucleotide-binding protein. The “core” of the TwoAYGGAY structure is shown, the Rfam (v11.0) model contains a further external stem that is not well conserved. Also, the reverse-complement (RevComp) of the Bacillaceae-1 is illustrated, this strand has the matches to the CsrA binding motif and the original discoverers of this ncRNA are not confident of the strand (personal communication, Weinberg Z).

### 3.2 Evolution

Non-coding RNAs are remarkably tolerant of genetic variation, as evident by the wide degree of sequence variation that can be found between evolutionarily related ncRNAs [66, 67, 68, 15]. However, structure frequently constrains the evolution of RNA sequences. That said, structures can also be dynamic. For example, motifs that confer structural stability can be exchanged over time, resulting in a rich and complex evolutionary history. This illustrates that studying the gain and loss of RNA motifs over evolutionary time-scales can help characterise the dynamic evolution of RNA sequences and structures.

**Figure 3.**
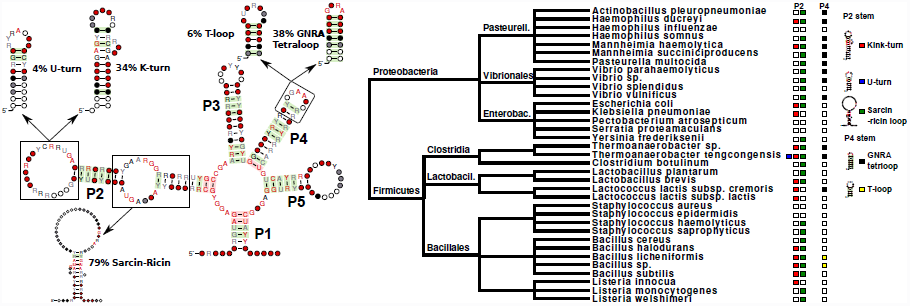
The Lysine riboswitch has substituted different motifs through its evolution. On the left is a representation of the consensus Lysine riboswitch secondary structure [61]. This has been annotated with the most frequent motifs the RMfam annotates in the Lysine Rfam (v11.0) seed alignment, the percentage of seed sequences hosting each motif is also indicated. On the right is an annotated species taxonomy that illustrates the phylogenetic nature of the motif distributions. We have also annotated each tip with the motifs hosted in the P2 and P4 stems. The red, blue, green, black and yellow boxes illustrate kink-turn, U-turn, sarcin-ricin loop, GNRA tetraloop and the T-loop, respectively.

A good example of this is the Lysine riboswitch. This is a convenient example, that for illustrative purposes that we will describe in further detail. As illustrated in Figure 3 many motifs may be exchanged, e.g. the U-turn motif with a k-turn in the P2 stem or the T-loop and the GNRA tetraloop in stem P4. Interestingly, the motif distributions are relatively clade-like, with closely related riboswitches more likely to share motifs, e.g. the GNRA tetraloop found in the Pasteurellales and Vibrionales taxonomic groups. This type of annotation information is valuable for researchers investigating the structure and evolution of RNA families.

### 3.3 Curation

Another use of the results presented in this work is of importance for the curators of RNA alignments and sequences [12, 69, 70]. Until now it has been difficult to analyse the evolutionary conservation of motifs in the context of an alignment, although some progress has been made when crystallographic data is available, e.g. the RNASTAR collection of structural RNA alignments [70]. With the help of RMfam, malformed alignments can be detected and corrected where conserved RNA motifs are incorrectly aligned. We illustrate an example of this for the Rfam (v11.0) 5S rRNA alignment that contains a misaligned, yet highly conserved sarcin-ricin motif (see Figure S45), and for the Rfam RsmY alignment, which is a CsrA binding sRNA. The RsmY alignment has a mis-annotated consensus structure that does not include a further CsrA binding motif (see Figure S46). These motifs generally occur in pairs, as CsrA is a homodimeric protein, with each half of the protein binding a motif [71, 72].

## 4. DISCUSSION & CONCLUSION

The chief motivation for this work is to functionally characterise novel ncRNAs. Our vision for the RMfam resource is to annotate RNAs of unknown function (e.g. [8]). These motif annotations will help develop further functional hypotheses and accelerate experimental characterisation.

In this work, we have shown that RMfam is surprisingly accurate. Despite the fact that the average RMfam motif consists of just 34.3 nucleotides and 10.9 basepairs, we show that the covariance models are specific enough to distinguish between motif-hosting sequences and negative control sequences (See Figures 1 and S10-S43). Our approach shows improved performance when evolutionary information encoded in Rfam sequence alignments is incorporated into the predictions. We hypothesise that annotated genome alignments may be a useful source of motifs and we will investigate this idea further in future. As a discovery tool this resource has already made some useful predictions. We have predicted the existence of two new CsrA binding ncRNAs, potentially the first of this class of regulatory molecules to be found outside of the Gammaproteobacteria. However, further work needs to be carried out to validate this claim.

### 4.1 Future work and potential applications

We have identified some future developments and applications for the RMfam resource. We plan to continue developing the accuracy of the motif annotation tools as well as increase the access to RMfam annotations via other databases and expand the number of motifs included in RMfam. Furthermore, it may be possible to boost the accuracy of RNA secondary structure prediction tools by constraining these with predicted motifs. We elaborate further on these ideas below.

The Lysine riboswitch example raises the possibility that certain types of motif are preferentially exchanged during the evolution of ncRNAs. Do stable hairpin motifs such as the GNRA and T-loops replace each other more frequently than we expect by chance? This would blur the lines between our understanding of homologous and analogous structures [73]. Another possibility is that certain motifs co-occur more frequently than we expect. For example, are k-turns more frequently closed by U-turns than we expect? If correct, these enrichments of favoured exchanges and co-occurances could be used to increase our confidence in motif annotations and can assist with the design of functional RNAs.

Typical RNA structure prediction methods to not incorporate information about RNA motifs. We propose that RMfam predictions can be used as constraints for existing RNA structure prediction software, thus improving the accuracy of structure prediction tools which can often be inaccurate [74]. This approach is analogous to the fragment-library approach that is frequently used for tertiary structure prediction [75].

Another application for RMfam covariance models is as a pre-filter to accelerate the more complex methods, for example, the Bayesian network approach implemented in RMdetect [19].

Increasing the access of motif annotations is another goal of the authors. We are active in the Rfam and RNAcentral consortia, both of which curate non-coding RNAs, the former ncRNA alignments and the latter ncRNA sequences [12, 69, 13]. Our results show that curators can benefit greatly from motif annotations (see Figures S44-S45) and it is likely that RMfam annotations will be incorporated into these databases in future releases.

New technologies such as the sequencing of cross-linked RNA and protein are a potential source of new RNA-protein motifs. In the future we will mine these datasets [76, 77, 78] for new additions to the RMfam database. Furthermore, we will continue to add new motifs to RMfam as they are published.

Finally, as previously mentioned, the specificity of the RMfam annotations is generally low. However, incorporating the genomic and taxonomic context of annotations into the predictions may result in performance gains. For example, Shine-Dalgarno and rho-independent terminators are generally located in bacterial sequences and at the extremities of annotated genes. A probabilistic incorporation of contextual information will likely result in further performance gains.

In summary, we have developed a resource for annotating diverse sets of RNA motifs in nucleotide sequences and alignments. We have proven the accuracy using benchmarks, and the utility of this resource for alignment curation, evolutionary analyses and shown that it has some promise for the prediction of RNA function.

## 5. ACKNOWLEDGEMENTS

This work would not be possible without input from a large community of RNA researchers that openly share their results. It has benefited from many discussions with members of the Xfam Consortium, the RNA Ontology Consortium and attendees of the 2012 Benasque Meeting on RNA. A special thanks to Lars Barquist, Elena Rivas, Rob Knight, Eric Westhof, Zasha Weinberg, Anthony Poole, Peter Fineran and two anonymous reviewers for their valuable contributions.

PPG is supported by a Rutherford Discovery Fellowship from Government funding, administered by the Royal Society of New Zealand.

